# Single-cell profiling of tumor-reactive CD4^+^ T-cells reveals unexpected transcriptomic diversity

**DOI:** 10.1101/543199

**Authors:** Assaf Magen, Jia Nie, Thomas Ciucci, Samira Tamoutounour, Yongmei Zhao, Monika Mehta, Bao Tran, Dorian B. McGavern, Sridhar Hannenhalli, Rémy Bosselut

## Abstract

Most current tumor immunotherapy strategies leverage cytotoxic CD8^+^ T cells. Despite evidence for clinical potential of CD4^+^ tumor-infiltrating lymphocytes (TILs), their functional diversity has limited our ability to harness their activity. To address this issue, we have used single-cell mRNA sequencing to analyze the response of CD4^+^ T cells specific for a defined recombinant tumor antigen, both in the tumor microenvironment and draining lymph nodes (dLN). Designing new computational approaches to characterize subpopulations, we identify TIL transcriptomic patterns strikingly distinct from those elicited by responses to infection, and dominated by diversity among T-bet-expressing T helper type 1 (Th1)-like cells. In contrast, the dLN response includes follicular helper (Tfh)-like cells but lacks Th1 cells. We identify a type I interferon-driven signature in Th1-like TILs, and show that it is found in human liver cancer and melanoma, in which it is negatively associated with response to checkpoint therapy. Our study unveils unsuspected differences between tumor and virus CD4^+^ T cell responses, and provides a proof-of-concept methodology to characterize tumor specific CD4^+^ T cell effector programs.
Targeting these programs should help improve immunotherapy strategies.

**One Sentence Summary:** Single-cell RNA sequencing reveals novel and highly diverse transcriptomic patterns characteristic of CD4^+^ T cell responses to tumors.

## Introduction

Immune responses have the potential to restrain cancer development, and most immunotherapy strategies aim to reinvigorate T cell function to unleash effective anti-tumor immune responses (*1–5*). Cytotoxic CD8^+^ T lymphocytes are being exploited in clinical settings due to their ability to recognize tumor neo-antigens and kill cancer cells (*3,6*). However, effective anti-tumor immunity relies on a complex interplay between diverse lymphocyte subsets that remain poorly characterized. CD4^+^ T helper cells, which are essential for effective immune responses and control the balance between inflammation and immunosuppression (*4,7–9*), have recently emerged as potential therapeutic targets (*4–6,10–14*). CD4^+^ helper cells contribute to the priming of CD8^+^ T cells and to B cell functions in lymphoid organs (*4,15,16*). CD4^+^ T helper type-1 (Th1) cells secrete the cytokine IFN-*γ* and affect tumor growth by targeting the tumor microenvironment (TME), antigen presentation through MHC class I and II, and other immune cells (*17–22*). Conversely, Th2 cells can promote tumor progression and regulatory T cells (Treg) mediate immune tolerance, suppressing the function of other immune cells and thus preventing ongoing anti-tumor immunity (*23–25*).

Despite the anti-tumor potential of CD4^+^ T cells, disentangling their functional diversity has been the limiting factor for pre-clinical and clinical progress. While several studies have assessed the transcriptome of Treg cells or their tumor reactivity (*25,26–31*), the functional diversity of conventional (non-Treg) tumor-infiltrating lymphocytes (TILs) has remained poorly understood. Population studies have limited power at identifying new, and especially rare functional cell states. Conventional single-cell approaches (e.g. flow or mass cytometry) overcome this obstacle but are necessarily restricted to hypothesis-based targets because of the number of parameters they can analyze. Furthermore, most previous studies, whether of human or in experimental tumors, did not distinguish tumor antigen-specific from bystander CD4^+^ T cells, even though bystanders may form the vast majority of conventional (non-Treg) T cells in the TME (*28,30–35*), in particular in draining lymphoid organs, where immune responses are typically initiated.

To address these challenges, we applied the resolution of single-cell RNA-sequencing (scRNAseq) to a tractable experimental system assessing tumor-specific responses both in the tumor and in lymphoid organs, and we designed new computational analyses to identify transcriptomic similarities. Our analyses dissect the complexity of the CD4^+^ T cell response to tumor antigens and identify broad transcriptomic divergences between anti-tumor and anti-viral responses. Emphasizing the power of this approach, new transcriptomic patterns identified in the present study are also found in CD4^+^ T cells infiltrating human tumors and correlate with response to checkpoint therapy in human melanoma.

## Results and Discussion

### Tracking tumor-specific CD4^+^ T cells

We set up a tractable experimental system to study tumor antigen-specific CD4^+^ T cells. We retrovirally expressed the lymphocytic choriomeningitis virus (LCMV) glycoprotein (GP) in colon adenocarcinoma MC38 cells, using a vector expressing mouse Thy1.1 as a reporter (**Figure S1A**). Subcutaneous injection of the resulting MC38-GP cells produced tumors allowing analysis of immune responses by day 15 after injection. We tracked GP-specific CD4^+^ T cells through their binding of tetramerized I-A^b^ MHC-II molecules associated with the GP-derived GP66 peptide (*36*). Such CD4^+^ cells were found in the tumor and draining lymph node (dLN) of MC38-GP tumor bearing mice, but neither in non-draining LN (nLN) from MC38-GP mice, nor in mice carrying control MC38 tumors (**Figure S1B**).

To study the CD4^+^ T cell response to tumor antigens, we aimed to produce genome-wide single cell mRNA expression profiles (scRNAseq) in CD4^+^ TILs and CD4^+^ dLN cells. We sorted GP66-specific T cells from dLNs, as these were the only dLN CD4^+^ T cells for which tumor specificity could be ascertained. Among TILs, we noted that ∼87% of GP66-specific CD4^+^ T cells expressed Programmed Cell Death 1 (PD-1, encoded by *Pdcd1,* **Figure S1C**), a marker of persistent antigenic stimulation (*37*). Thus, to obtain a broad representation of antigen-specific TILs, not limited to GP-specific cells, we used PD-1 expression as a surrogate for tumor antigen specificity and purified tumor CD4^+^ CD44^hi^ PD-1^+^ T cells (PD-1^hi^ TIL) for scRNAseq. We verified critical conclusions of the scRNAseq analyses by flow cytometry, comparing GP66-specific and PD-1^hi^ TILs.

### Tumor-responsive CD4^+^ T cells are highly diverse

We captured GP66-specific dLN and PD-1^hi^ TIL (dLN and TILs hereafter, respectively) CD4^+^ cells using the 10x Chromium scRNAseq technology (*38*); additionally, we captured GP66-specific spleen CD4^+^ T cells from LCMV (Armstrong strain)-infected mice (*36*) as a technical and biological reference (**Figure S1D**, called ‘LCMV cells’ here). We excluded cells of low sequencing quality (low number of detected genes), potential doublets, and B cell contaminants, leaving 566 dLN, 730 TIL, and 2163 LCMV CD4^+^ T cells for further analyses (**Table S1**).

We defined groups of cells sharing similar transcriptomic profiles using Phenograph clustering (*39*). Consistent with previous studies (*40*), LCMV cells segregated into follicular helper (Tfh, providing help to B cells) and type-1 helper (Th1, secreting the cytokine IFN-*γ*) T cells, among other subsets (**Figure S2A**). Tfh cells expressed *Tcf7* (encoding the transcription factor Tcf1), *Cxcr5,* and *Bcl6*, whereas Th1 cells expressed *Tbx21* (encoding the transcription factor T-bet), *Ifng* (IFN-*γ*), and *Cxcr6*. Low resolution clustering identified 5 groups of TILs and dLN cells (**Figure S2B**). Groups I and II had features of Th1 cells, although group II differed by higher expression of the chemokine receptor *Cxcr3* and lower expression of *Ifng*. Group III expressed genes typical of Treg cells, including *Foxp3* and *Il2ra*, encoding CD25 (IL-2R*α*). Groups IV and V expressed Tfh cell genes, including *Bcl6* and *Cxcr5*, and group IV *Ccr7*, which preferentially marks memory cell precursors at the early phase of the immune response (*40,41*).

To further dissect these populations, we developed a user-independent, data-driven approach to increase clustering resolution while controlling for false discovery. Applying such high-resolution clustering separately to TILs and dLN cells, we identified 15 clusters (TIL clusters t1-t7 and dLN clusters n1-n8), refining the original five main groups (**Figure 1A**). Revealing unexpected diversity among Th1-like TILs, group I and II resolved into 5 subpopulations, including a distinct cluster (t5) expressing higher levels of *Il7r* (encoding IL-7R*α*) and lower levels of *Tbx21* and *Ifng*. Only cluster group III (Tregs) included both TIL and dLN cells, which expressed variable levels of *Tbx21*. Groups IV and V, the bulk of dLN cells, resolved into 5 and 2 clusters, respectively. Consistent with flow cytometric analysis, dLN cells neither expressed high levels of T-bet, the product of *Tbx21*, nor exhibited Th1 attributes; in contrast, most TILs expressed T-bet, even if at various levels (**Figure 1A and S2C, D**).

**Fig. 1:**
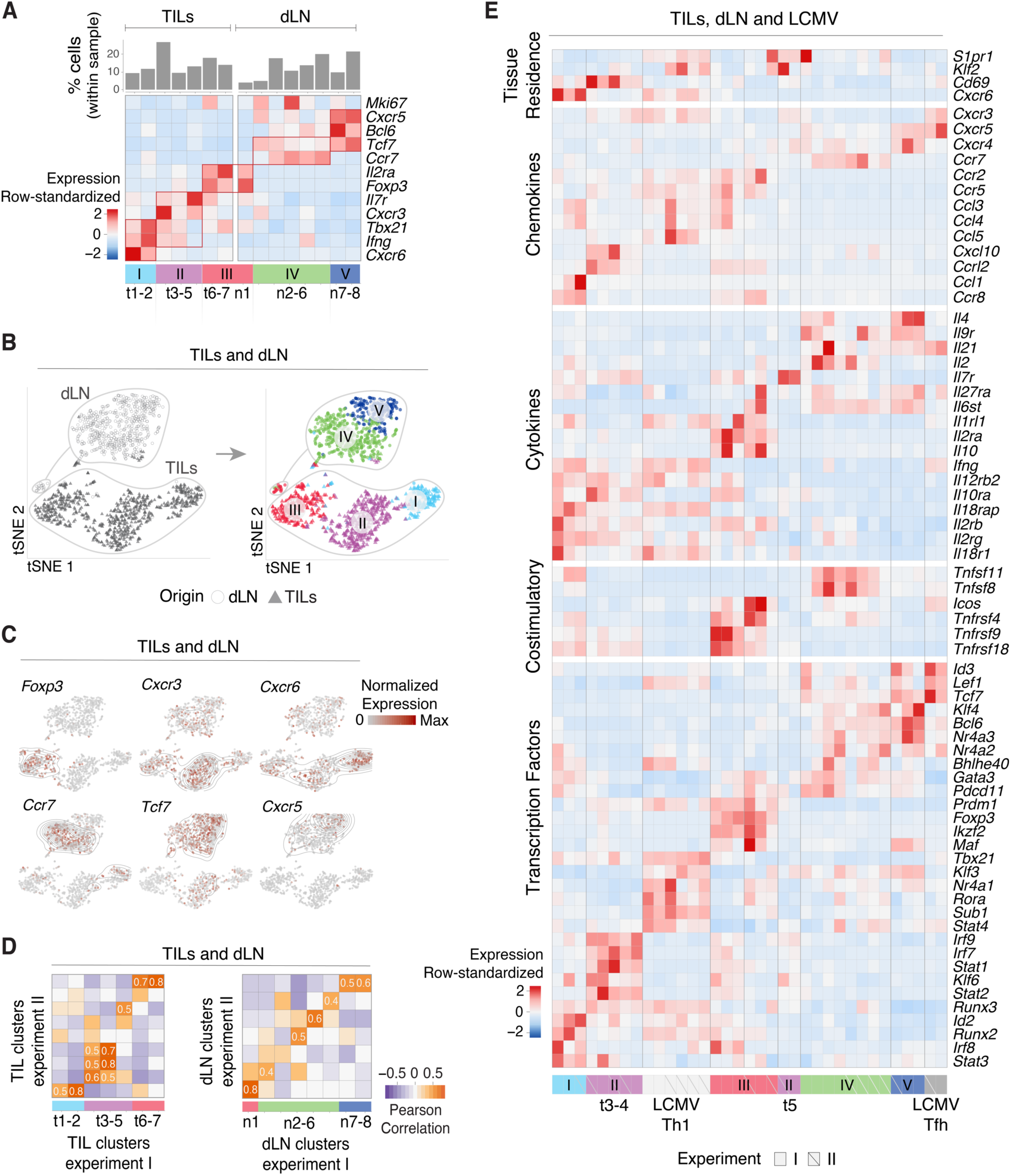
Characterization of CD4^+^ TIL, dLN and LCMV transcriptomes by scRNAseq. **(A-D)** TILs and dLN cells from WT mice at day 14 post MC38-GP injection analyzed by scRNAseq. **(A)** Heatmap shows row-standardized expression of selected genes across TIL and dLN clusters. Bar plot indicates the number of cells in each cluster relative to the total TIL or dLN cell number. **(B)** tSNE display of TILs and dLN cells, grey-shaded by tissue origin (**left**) or color-coded by main group (**right**, as defined in A). **(C)** tSNE (TIL and dLN cell positioning as shown in B) display of normalized expression levels of selected genes. **(D)** Heatmap shows Pearson correlation between clusters’ FC vectors (as defined in text) across the two replicate experiments for TILs (**left**) and dLN (**right**). **(E)** TILs, dLN and LCMV cells from replicate experiments I and II analyzed by scRNAseq. Heatmap shows row-standardized expression of selected genes across clusters. Group II (purple) t5 separated into a distinct component from t3-4 (as defined in text).

To support these observations, we analyzed pooled TILs and dLN cells by t-Distributed Stochastic Neighbor Embedding (t-SNE), a dimensionality reduction approach that positions cells on a two-dimensional grid based on transcriptomic similarity (*42*). Although performed on the pooled populations, t-SNE recapitulated the minimal overlap between TIL and dLN transcriptomic patterns (**Figure 1B, left**), irrespective of parameter selection (**Figure S2E**) and even after controlling for potential confounders (**Figure S2F and Supplementary Note and Figure**). Remarkably, cluster groups I-V almost completely segregated from each other when projected on the t-SNE plot (**Figure 1B, right**). Overlay of gene expression confirmed co-localization of cells expressing cluster-characteristic genes (**Figure 1C**).

To verify the robustness of these observations, we analyzed a biological replicate consisting of 1123 TILs and 675 dLN GP66-specific cells captured from a separate set of tumors (**Figure S2G and Table S1**). Because batch-specific effects can confound co-clustering from distinct experiments, we separately clustered cells from each replicate. To compare these clusters, we evaluated the correlation between cluster-specific fold-change (FC) vectors, defined internally to each replicate, that recorded expression of each gene in a cluster relative to all other clusters in that replicate. We found significant inter-replicate matches for most clusters (**Figure 1D**), supporting the reproducibility of the underlying transcriptomic patterns. Thus, scRNAseq analysis of tumor-specific CD4^+^ T cells identifies an unsuspected diversity of transcriptomic programs in the TME and dLN.

### Correlation analyses mitigate tissue-context-specific factors

Comparison of TILs, dLN, and LCMV cells showed little overlap, including between TILs and dLN cells (**Figure S2H, left**). Thus, we considered that the impact of tissue of origin could be the primary driver of clustering and mask commonalities in effector programs. Indeed, most TIL subpopulations had attributes of tissue residency, including low *S1pr1* and *Klf2* expression, and high expression of *Cd69*, contrasting with LCMV and most tumor dLN clusters (**Figure 1E)** (*43*). Only group III Tregs, and separately cells undergoing cell cycle, clustered together regardless of origin (**Figure S2H, right**). This prompted us to search for potential underlying similarities among these disparate transcriptomic patterns. We found that data integration approaches designed to uncover similarities across experimental conditions could not overcome the separation resulting from biological context (**Figure S3A**), and could miss functionally relevant differences (e.g. between Foxp3^+^ and Foxp3^−^ TILs, **Figure S3B**) (*44*). Thus, we considered the correlation analysis used above for cluster matching. This analysis distributed the 40 reproducible clusters (out of a total of 47 from all experiments) into 6 ‘meta-clusters’ (with manual curation attaching meta-cluster 1b to 1a), of which four (meta-clusters 1, 3, 5 and 6) comprised cells of more than one tissue context (**Figure 2A and Table S2**). Thus, the correlation analysis establishes relatedness among transcriptomic patterns identified by conventional clustering.

**Fig. 2:**
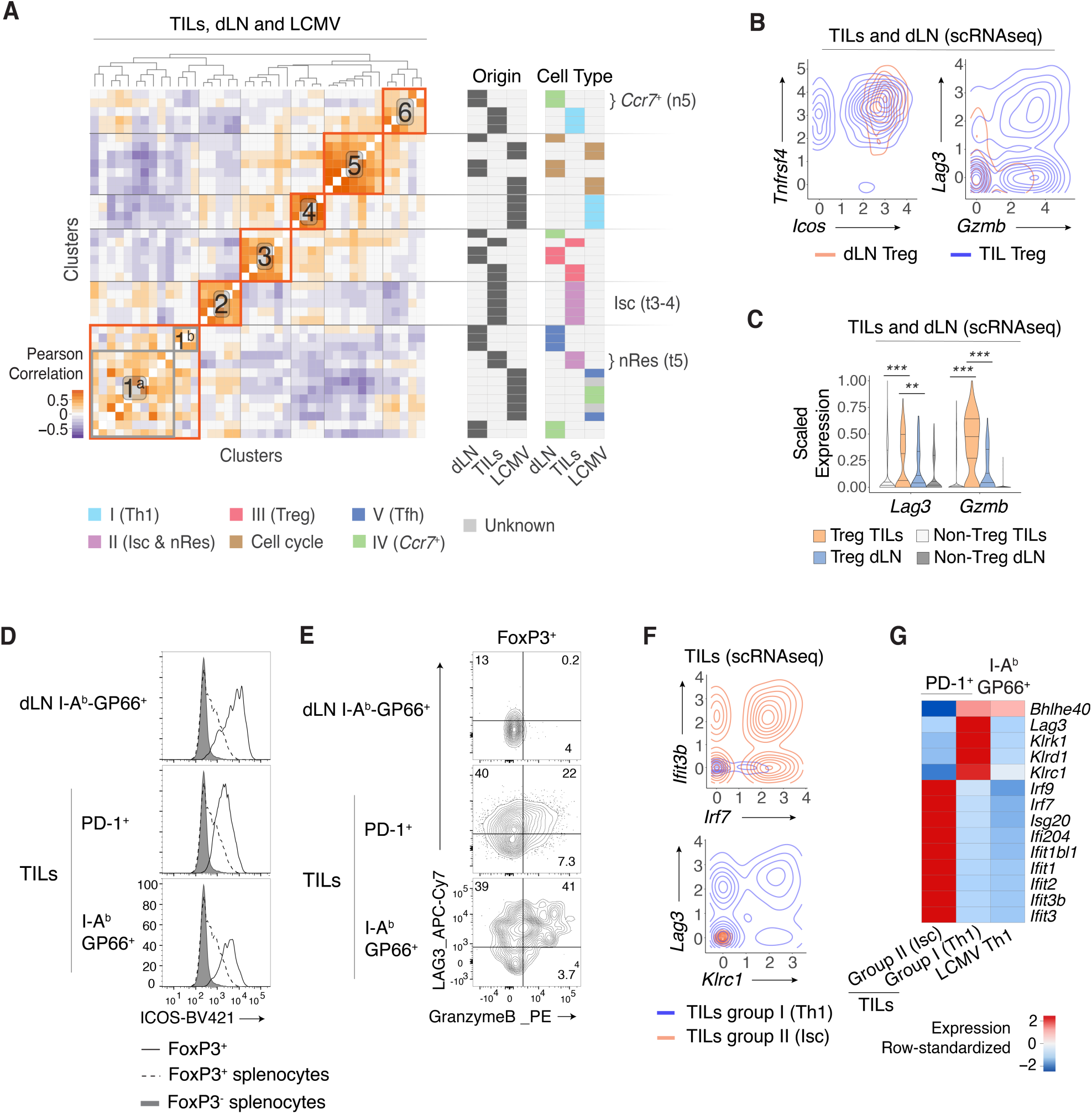
Treg and Th1-like transcriptomic patterns. **(A)** Heatmap defines meta-clusters based on Pearson correlation between TIL, dLN and LCMV cluster FC vectors (as defined in text) (**left**). Indicator tables show tissue origin and cell type color-code per cluster (**right**). **(B-E)** Comparison of dLN Tregs and TIL Tregs (respectively clusters t6-7 and n1 as shown in Fig. 1A). **(B)** Contour plots of dLN Treg (orange) or TIL Treg (blue) cell distribution according to scRNAseq-detected normalized expression of *Icos* vs. *Tnfrsf4* (**left**) and *Gzmb* vs. *Lag3* (**right**). **(C)** Violin plot of *Lag3* and *Gzmb* scRNAseq expression in Treg vs. non-Treg TIL and dLN populations (Unpaired T test, * p < 0.01, ** p < 0.001); bands indicate quartiles (25^th^, 50^th^ and 75^th^ quantile). **(D)** Overlaid flow cytometry expression of ICOS in Foxp3^+^ TILs and dLN cells and Foxp3^+^ or Foxp3^-^ CD4^+^ splenocytes from tumor-free control mice. **(E)** Flow cytometry contour plots of Granzyme B vs. LAG3 in Foxp3^+^ TILs and Foxp3^+^ dLN cells. **(F-G)** Comparison of TIL Th1 and Isc (respectively clusters t1-2 and t3-4 as shown in Fig. 1A) to LCMV Th1 (as shown in Fig. 1E and S2A) **(F)** Contour plots of Th1 (orange) and Isc (blue) TIL distribution according to scRNAseq-detected normalized expression of *Irf7* vs. *Ifit3b* (**top**) and *Klrc1* vs. *Lag3* (**bottom**). **(G)** Heatmap shows row-standardized expression of differentially expressed genes across TILs group II Isc, TILs group I Th1 and LCMV Th1.

### Characterizing transcriptomic similarities

We further characterized the meta-clusters by identifying their defining overexpressed genes. In addition to *Foxp3* and *Il2ra*, genes driving meta-cluster 3 (Treg, group III) included *Ikzf2, Tnfrsf4*, encoding Ox40, and *Ico*s, which we verified by flow cytometry (**Figures 1E, 2B left, and 2D**). In contrast, *Gzmb* (encoding the cytotoxic molecule Granzyme B) and *Lag3* were overexpressed in TIL Tregs relative to dLN Tregs (and to other TIL subsets) (**Figure 2B right, C, E**). Thus, the similarity analysis both confirmed the shared Treg circuitry across TILs and dLN and identified TIL-specific *Gzmb* cytotoxic gene expression in TIL Tregs.

Contrasting with Treg clusters, the correlation analysis failed to detect similarities between the three groups of T-bet-expressing cells. These cells, which showed heterogeneous *Tbx21* levels, were distributed into meta-clusters 2 (TILs group II, t3-4), 4 (LCMV cells) and 6 (TILs group I, t1-2) (**Figure 2A**). The two TIL meta-clusters showed multiple differences from LCMV-responsive Th1 cells, including higher expression of *Il12rb, Il7r* and *Il10ra*, and distinct patterns of transcription factor, chemokine and chemokine receptor expression. Relative to the other T-bet-expressing cells, TILs group II (t3-4) differed by high expression of multiple type I IFN-induced genes, including transcription factors *Irf7* and *Irf9* (**Figure 2F top, 2G, S3C**). Co-expression of these genes with T-bet was unexpected, as T-bet normally repress genes induced by type I IFN (*68*). We designated group II t3-4 as interferon stimulated clusters (Isc). Group I t1-2 TIL clusters (Th1 hereafter) specifically expressed *Lag3* and Killer Cell Lectin (Klr) genes (**Figure 2F bottom, 2G, S3C**), characteristic of terminally differentiated effector cells (*45*). Flow cytometry verified that Th1 TILs did not express the Natural Killer (NK) T cell-specific transcription factor PLZF, indicating they were not NK T cells (**Figure S3D**). Compared to Isc, Th1 clusters had higher expression of *Bhlhe40*, a transcription factor controlling inflammatory Th1 fate determination (*46,47*). A recent study of human colon cancer identified a CD4^+^ TIL Th1 subset with elevated *Bhlhe40* expression (*31*). This subset is clonally expanded and enriched in tumors with micro-satellite instability, suggesting specificity for tumor antigens. The mouse Th1 TILs identified in our study had higher expression of 40 genes from the human colon TIL Th1 signature, including *Bhlhe40* and *Lag3* (**Table S3**), with a significant (p=0.001) skewing towards this signature detected by GSEA (*48*). However, mouse Th1 TILs lacked expression of other components of the human signature, including *Gzmb* and *Irf7*, suggesting that the impact of *Bhlhe40* expression on TIL transcriptomes is in part context-specific.

Meta-cluster 6 unexpectedly associated Th1 TILs and a dLN *Ccr7*^+^ cluster (Group IV cluster n5) (**Figure 2A**), suggesting a potential link between TILs and dLN. The association was driven by transcriptional regulators *Bhlhe40* and *Id2,* and TNF superfamily members *Tnfsf8* (encoding CD30L) and *Tnfsf11* (RANKL) (**Figures 3A and 1E**). The potential connection between *Ccr7*^+^ dLN cells and Th1 TILs was specific to *Ccr7*^+^ cluster n5, which segregated from n6 and other dLN subsets (Tfh and Treg) based on higher expression of *Ifng* (but not *Tbx21*) and *Cd200* (**Figure 3B**). Flow cytometry identified a corresponding CD200^hi^ subset among Cxcr5^lo^ Ccr7^+^ but not Cxcr5^+^ Ccr7^−^ (Tfh) GP66-specific cells (**Figure 3C, S3E and S3F**). dLN *Ccr7*^+^ clusters t5-6 shared features with central memory precursor CD4^+^ T cells (Tcmp) identified in LCMV infection (*40*) (**Table S3 and Figure 1E**). This includes expression of *Tcf7*, a transcription factor important to prevent T cell terminal differentiation and for CD8^+^ T cells responsiveness to PD-1 blockade (*49–56*). However, the correspondence between MC38-GP dLN *Ccr7*^+^ clusters and the LCMV Tcmp signature was only partial (**Table S3**).

**Fig. 3:**
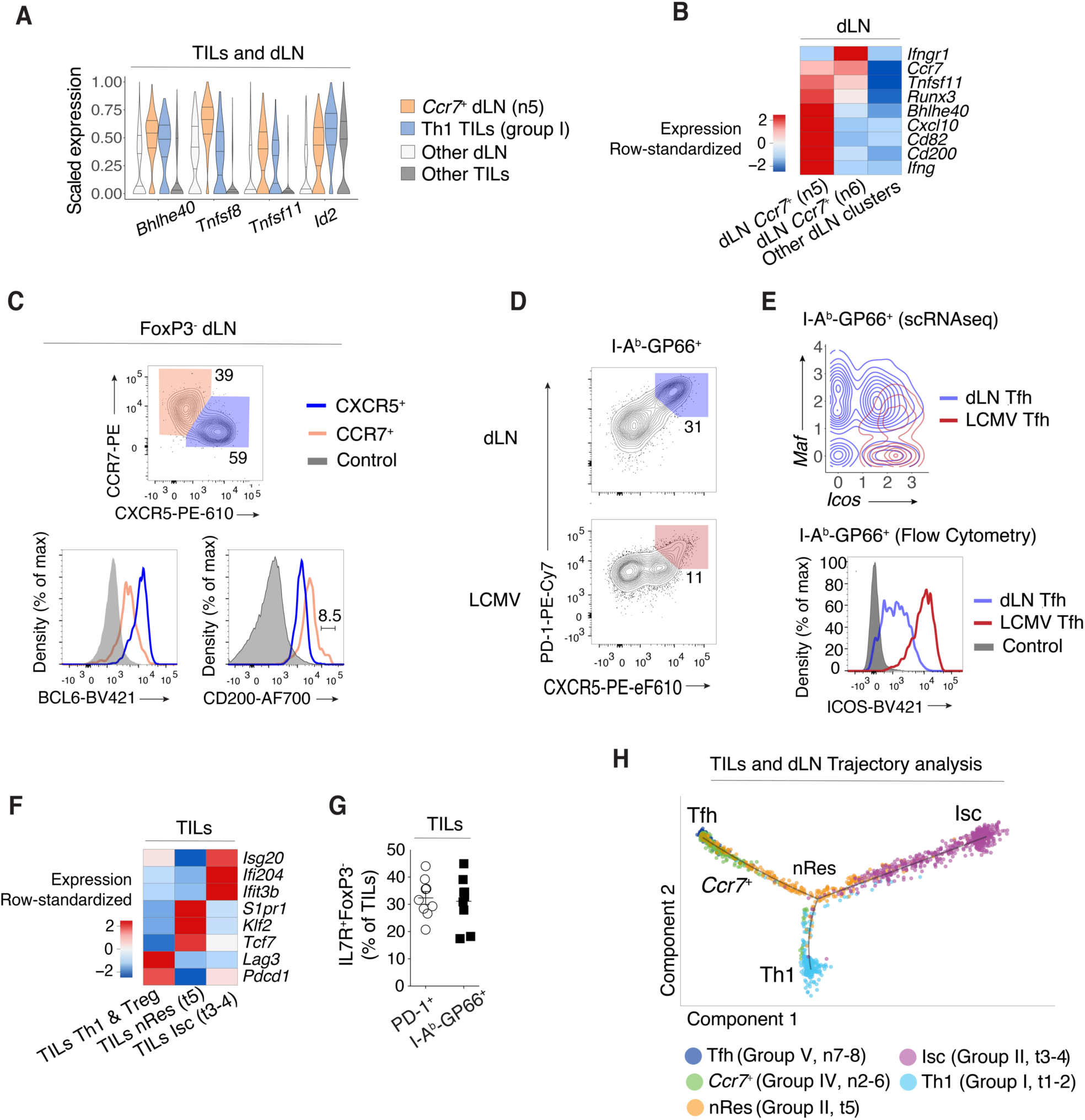
Transcriptomic continuum between TIL and dLN tumor-reactive cells. **(A)** Violin plots of differentially expressed genes across TILs group I Th1, dLN group IV *Ccr7*^+^ (respectively clusters t1-2 and n5 as shown in Fig. 1A) and all other TIL and dLN populations. **(B)** Heatmap shows row-standardized expression of differentially expressed genes across dLN *Ccr7*^+^ clusters (group IV n5-6) and other dLN clusters (Treg and Tfh clusters n1 and n7-8, respectively). **(C)** Top panel shows flow cytometry contour plots of CXCR5 vs. CCR7 in Foxp3^-^ dLN cells. Bottom panel shows overlaid protein expression of BCL6 and CD200 in CCR7^+^ and CXCR5^+^ dLN cells and naive CD4^+^ splenocytes from tumor-free control mice. **(D)** Flow cytometry contour plots of CXCR5 vs. PD-1 in dLN and LCMV cells. **(E)** Contour plot of dLN (red, clusters n7-8) and LCMV (blue) Tfh cell distribution according to scRNAseq-detected normalized expression of *Icos* vs. *Maf* (**top**). Overlaid protein expression of ICOS in dLN and LCMV PD-1^+^CXCR5^+^ (Tfh) cells and naive CD4^+^ splenocytes from tumor-free control mice (**bottom**). **(F)** Heatmap shows row-standardized expression of differentially expressed genes across TILs Isc and nRes clusters (as defined in text, group II t3-4 and t5, respectively) and all other TIL clusters (Th1 and Treg clusters t1-2 and t6-7, respectively). **(G)** Fractions of IL7R^+^Foxp3^-^ cells out of total PD-1^+^ or GP66^+^ TILs. **(H)** Trajectory analysis of PD-1^+^ TILs and GP66^+^ dLN cells indicating individual cells assignment into a transcriptional continuum trajectory. nRes cluster (t5) is color-coded in orange in contrast to annotations in other figures.

Meta-cluster 1 comprised LCMV Tfh clusters and dLN group V Tfh clusters (**Figure 2A**). We verified that the abundance of dLN Tfh cells was similar in mice carrying MC38-GP and MC38 tumors (**Figure S3G**), indicating that this response is not a consequence of GP expression. Flow cytometric analysis confirmed key Tfh attributes in dLN and LCMV cells (**Figure 3D**), although dLN Tfh cells differed from LCMV-responsive Tfh cells by lower expression of *Icos* and the upregulation of the transcription factor *Maf* (**Figure 3E, 1E and S3H**). Unexpectedly, meta-cluster 1 associated the dLN and LCMV Tfh clusters with TIL group II cluster t5, characterized by *Il7r* expression (**Figures 2A and 1A**), based in part on intermediate expression of *Tcf7* (1.6 fold relative to other TIL subpopulations) (**Figure 3F and 1E**). Flow cytometric analysis confirmed the abundance of GP66-specific IL-7R^+^ TILs (**Figure 3G**). In addition, the *Tcf7*^int^ t5 cluster showed expression of the transcription factor *Klf2* and its downstream target Sphingosine-1-phosphate receptor 1 (*S1pr1*). This indicated retention of a cell trafficking transcriptional program (*57*) (**Figure 3F and 1E**) and contrasted with the interferon-driven Isc TILs. Thus, we designated cluster t5 of group II TILs as putative non-resident cells (nRes hereafter).

To further delineate the relationships between cell clusters, we used Reversed Graph Embedding (*58*), which has been used to estimate progression through transcriptomic states. This placed the dLN Tfh and TIL Th1 and Isc at the end of an inferred path (**Figure 3H**), nRes TILs in the middle of the continuum and *Ccr7*^+^ dLN cells between Tfh and nRes. These analyses, combined with the similarities described by meta-clustering, support the notion that the tumor-responsive CD4^+^ T cell response may be characterized as a transcriptomic continuum; they confirm the transcriptomic distance between Th1 and Isc TILs, even though both subsets express T-bet, the Th1-defining factor.

### TILs subpopulation-specific dysfunction gene programs

We reasoned that expression of a dysfunction-exhaustion program (*59,60*) may account for the limited relatedness between LCMV and TIL Th1 cells, as TILs expressed multiple exhaustion marks (**Figure 4A**), and were sorted for PD-1 expression for scRNAseq. To assess the impact of exhaustion on TIL subpopulation, we defined TIL Th1, Isc, nRes and Treg gene signatures as the genes preferentially expressed in each subpopulation relative to all other TILs (**Table S4**). We found a significant overlap between multiple viral-response exhaustion gene signatures (MSigDB) (*61*) and the Th1 and Treg signatures (**Table S5**). Separate analysis of a previously reported gene signature characterizing CD4^+^ T cell dysfunction during chronic infection (*62*) indicated a significant overlap with the Isc signature, but not with Th1 and Treg signatures (**Figure S4A, Table S6**).

**Fig. 4:**
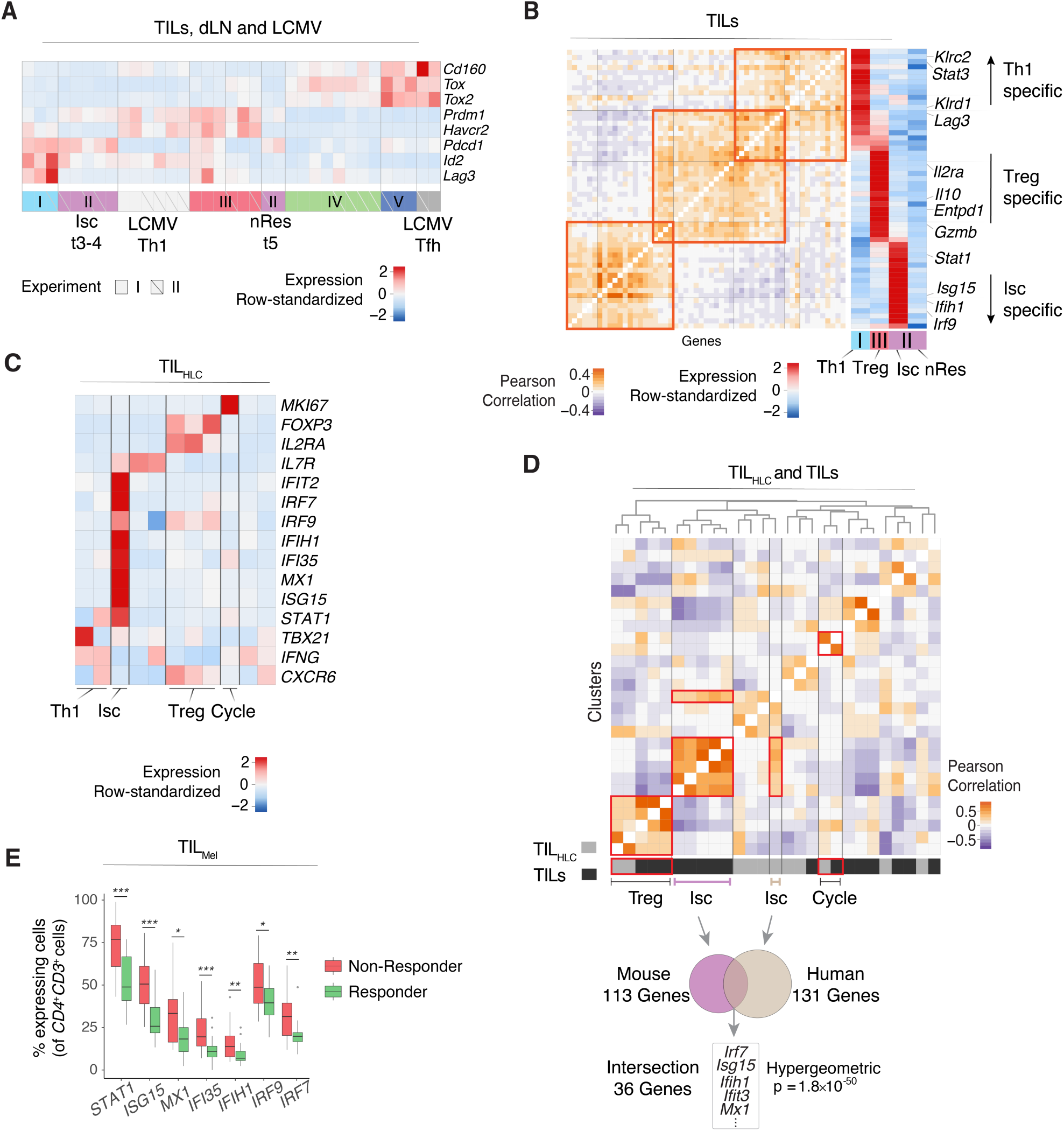
Correspondence to human data and dysfunction gene signatures. **(A)** Heatmap shows row-standardized expression of selected exhaustion genes across TIL, dLN and LCMV clusters from replicate experiments I and II. **(B)** Analysis of IL-27 signature genes overlapping with TIL subpopulation characteristic genes. Heatmap shows Pearson correlation (**left**) and row-standardized expression of overlapping genes across TIL Th1, Treg, Isc and nRes cells (respectively clusters t1-2, t6-7, t3-4 and t5 as shown in Fig. 1A) (**right**). **(C)** Analysis of human liver cancer TIL_HLC_. Heatmap shows row-standardized expression of selected genes across TIL_HLC_ clusters. **(D)** Heatmap defines meta-clusters based on Pearson correlation between TIL_HLC_ and MC38-GP TIL clusters (**top**). Overlap of genes characteristic of human liver TIL Isc cluster with mouse TIL Isc gene signature (**bottom**). **(E)** Analysis of human melanoma TIL_Mel_. Box plots show the percentage of cells expressing selected interferon signaling characteristic genes in *CD4*^+^*CD3*^+^ cells across responding and non-responding lesions (Unpaired Wilcoxon test, * p < 0.05, * p < 0.01, ** p < 0.001).

The latter result suggested heterogeneous expression of exhaustion genes among TIL subsets. We tested this possibility using a broader set of exhaustion genes shared across cancer and chronic infection (*63*). 55 genes from this set were also part of TIL Th1, Isc, or Treg signatures. However, their overlap was heterogeneous, identifying dysfunction programs specific of TIL subpopulations (**Figure 4B, Table S6**). Of note, we did not detect overlap between any dysfunction-exhaustion signature and nRes TILs (**Figure 4B, Table S6**). This is in line with these cells’ residual expression of *Tcf7*, which in CD8^+^ T cells marks cells with conserved responsiveness (*52–54,64*).

### The Isc IFN signature correlates with poor clinical prognosis in human tumors

Last, we examined if MC38-GP TIL transcriptomic patterns were observed in human tumors. We analyzed published CD4^+^ Human liver cancer TILs (TIL_HLC_) scRNAseq data pooled across six treatment-naive patients (*28*). High resolution clustering separated the TIL_HLC_ cells into 11 clusters, which could be combined into groups displaying features of Th1, Isc, Treg TILs and cells undergoing cell cycle (**Figure 4C**). While pooled analysis of CD4^+^ PD-1^+^ TILs from MC38-GP tumors (TIL) with TIL_HLC_ only identified similarities between cells undergoing cell cycle (**Figure S4B and S4C**), cluster correlation analysis indicated significant similarities between Tregs, cell cycle, and Isc clusters from TIL vs. TIL_HLC_ (**Figure 4D, top**). We focused on the Isc pattern, which differed the most from previously reported Th1 and Treg transcriptomic profiles. We found a significant overlap of overexpression patterns between TIL Isc and their human counterpart, including type I IFN-induced genes and *Irf7* (*65*) (**Figure 4D, bottom and Table S7**). Thus, the Isc signature newly identified among mouse CD4^+^ TILs is found in human tumors.

These finding were not unique to liver tumors, as analysis of CD4^+^CD3^+^ human melanoma TILs across 48 lesions (TIL_Mel_) (*33*) identified a cluster enriched in Isc characteristic genes, among other populations (**Figure S4D**). To investigate the relationships between Isc transcriptomic program and clinical prognosis, we evaluated the association between the expression in TIL_Mel_ of Isc signature genes (defined in MC38-GP TILs) and patient response to checkpoint therapy. Relative to responders, non-responsive tumors had significantly higher fractions of cells expressing Isc signature genes (49 out of 108 genes, adjusted p-value < 0.05), including *Stat1, Irf7* and *Irf9* (**Figure 4E and Table S8**). This indicated negative association between the Isc transcriptomic program and patient response to checkpoint therapy. Thus, the methods used in the present study identify transcriptomic programs shared by multiple tumor types and of potential prognostic significance.

In summary, using scRNAseq and data-driven computational approaches, the present study identifies an unsuspected diversity among tumor-responding CD4^+^ T cells. While recent scRNAseq studies had shed light on the Treg component of CD4^+^ TILs (*28,30–32*), our study assessed the transcriptomic patterns of both regulatory and conventional components, in the tumor itself and in draining lymphoid organs. We identify new transcriptomic patterns and find a heterogeneous distribution of exhaustion gene signatures among TILs subtypes, highlighting the need for extensive analyses of cell-specific effects of treatments targeting exhaustion genes.

Even though most conventional (Foxp3^−^) tumor-responsive TILs express T-bet, the Th1-defining transcriptional regulator, our study identifies novel and diverse transcriptomic patterns with unexpectedly little similarity to prototypical virus-responsive Th1 cells. Thus, conventional helper effector definitions, derived from studies of responses to infection, are inaccurate descriptors of responses to tumors. The newly identified Th1-like transcriptome with marks of type I IFN stimulation, a driver of inflammation and immunosuppression in cancer (*66*), highlights this conclusion: it was observed among TILs but not LCMV-responding cells, even though LCMV drives a strong type I IFN innate immune response (*67*). Our cluster similarity analysis projects this interferon-responsive transcriptomic pattern onto human tumors, overcoming potential sample disparity, and demonstrates its association with response to checkpoint therapy.

Investigating tumor-specific T cell responses in draining lymphoid organs revealed striking differences with TILs. The absence of Th1 cells from tumor dLN was unexpected and contrasted with infections, including with LCMV or with *Leishmania major*, a typical Th1-driving parasite with kinetics of clinical progression similar to that of experimental tumors, and in which Th1 dLN cells are important contributors to the response (*69*). In contrast, the tumor elicited strong, tumor-specific Foxp3-negative Tfh-like responses in dLN. While Tfh differentiation may divert T cells from more efficient (e.g. IFN*γ*-producing) anti-tumor differentiation, it provides support for the tantalizing possibility that tumor-elicited B cell responses could be exploited against cancer (*70*). It is also possible that this subset includes a stem cell-like component similar to the Cxcr5^+^ CD8^+^ dLN T cells that serve as targets for immunotherapy targeting PD-1 signaling (*52*), or cells with similar properties in the tumor micro-environment (*54*).

In conclusion, this study provides a high-resolution characterization of tumor-reactive CD4^+^ T cell responses in lymphoid organs and the tumor microenvironment. We identify previously unrecognized transcriptomic patterns among tumor-specific T cells and provide an extensive mapping of the CD4^+^ T cell immune response against cancer. We describe new analytical approaches of broad applicability, including to clinical data, that combine high resolution dissection of transcriptomic patterns and synthetic data integration to identify correspondences between apparently unrelated cell differentiation states.

## Materials and Methods

### Mice

C57BL/6 mice were purchased from the National Cancer Institute Animal Production Facility and were housed in specific pathogen-free facilities. Animal procedures were approved by the NCI Animal Care and Use Committee.

### Cell lines and constructs

MC38 murine colon cancer cell lines (*71*) were obtained from Jack Greiner’s lab and cultured in DMEM that contained 10% heat-inactivated FCS, 0.1 mM nonessential amino acids, 1 mM sodium pyruvate, 0.292mg/ml L-glutamine, 100 pg/ml streptomycin, 100 U/mL penicillin, 10mM Hepes. MC38-GP cells were generated as follows: LCMV-*gp* gene was amplified from pHCMV-LCMV-Arm53b (addgene#15796) and inserted into pMRX-IRES-Thy1.1 by BamH1 and Not1. Then pMRX-Thy1.1 contained LCMV-*gp* gene was transfected into Plat E cell to package retrovirus. MC38 cell line was transduced by above retrovirus collection and followed by single cell sorting in 96-well plate after 48hs. The monoclonal cell lines were identified by flow cytometry and western blot.

### LCMV infection model and Tumor model

2 x 10^5^ pfu of LCMV Armstrong (*36*) were injected intra-peritoneal in 6-12 weeks old C57BL/6 mice. Mice were analyzed 7 days post infection. MC38 and MC38-GP tumor cells (0.5 × 10^6^) were subcutaneously injected into the flank of C57BL/6 mice.

### Antibodies

Antibodies for the following specificities were purchased either from Affymetrix Becton-Dickinson Pharmingen or ThermoFisher Ebiosciences: CD4 (RM4.4 or GK1.5), CD8β (H35-17.2), CD45.2 (104), CD45 (30-F11), TCRβ (H57-597), CD5 (53-7.3), B220 (RA3-6B2), Siglec F (E50-2440), NK1.1 (PK136), CD11b (M1/70), CD11c (N418), CD44 (356 IM7), IL7R (A7R34), CCR7 (4B12), CXCR5 (SPRCL5), Bcl6 (K112-91), Lag3 (C9B7W), Cxcr6(SA051D1), CD25(PC61.5), CD278(7E,17G9), PD-1 (J43), Foxp3(FJK-16s), Granzyme B(FGB12), Tbet (4B10), CD200(OX-90). Streptavidin, MHC tetramers loaded with the *Toxoplasma gondii* AS15 (*72*) and LCMV GP66 peptides (AVEIHRPVPGTAPPS and DIYKGVYQFKSV, respectively) were obtained from the NIH Tetramer Core Facility.

### Cell preparation and flow cytometry

Lymph node and spleen were prepared and stained as previously described (*73*). For TIL preparation, tumors were dissected 14 to 18 days post-injection, washed in HBSS, cut into small pieces, and subjected to enzymatic digestion with 0.25mg/ml liberase (Roche) and 0.5mg/ml DNAase I (SIGMA) for 30 minutes at 37 degrees. The resulting material were passed through 70um filters and pelleted by centrifugation at 1500rpm. Cell pellets were resuspended in 44% Percoll (GE Healthcare) on an underlay of 67% Percoll, and centrifuged for 20min at 1600 rpm without brake. TILs were isolated from the 44%/67% Percoll interface. Following isolation, cells were blocked with anti-FcγRIII/FcγRII (unconjugated, 2.4G2) and subsequently stained for flow cytometry. Staining for AS15:I-A^b^ tetramer, GP66:I-A^b^ tetramer and CXCR5 was performed at 37 degrees for 1 hour prior to staining for other cell surface markers. For intracellular staining, cell surface staining were preformed first, following fixation using the Foxp3-staining kit (eBioscience). Flow cytometry data was acquired on LSR Fortessa cytometers (BD Biosciences) and analyzed with FlowJo (TreeStar) software. Dead cells and doublets were excluded by LiveDead staining (Invitrogen) and forward scatter height by width gating. Purification of lymphocytes by cell sorting was performed on a FACS Aria or FACS Fusion (BD Biosciences).

### Single cell RNAseq

3000-13000 T cells sorted from LCMV infected or tumor-bearing mice were loaded on the Chromium platform (10X Genomics) and libraries were constructed with a Single Cell 3′ Reagent Kit V2 according to the manufacturer instructions. Libraries were sequenced on multiple runs of Illumina NextSeq using paired-end 26×98bp or 26×57bp to reach a sequencing saturation greater than 70% resulting in at least 49000 reads/cell.

### scRNA-seq data pre-processing

De-multiplexing, alignment to the mm10 transcriptome and unique molecular identifier (UMI) calculation were performed using the 10X Genomics Cellranger toolkit (v2.0.1, http://software.10xgenomics.com/single-cell/overview/welcome). Pre-processing, dimensionality reduction and clustering analyses procedures were applied to each dataset (that is, specific tissue origin in each experiment) independently to account for dataset-specific technical variation such as sequencing depth and biological variation in population composition, as follows. We filtered out low quality cells with fewer than 500 detected genes (those with at least one mapped read in the cell). Potential doublets were defined as cells with number of detected genes or number of UMIs above the 98^th^ quantile (top 2% owing to up to 2% estimated doublets rate in the 10X Chromium system). Potentially senescent cells (more than 10% of the reads in the cell mapped to 13 mitochondrial genes) were also excluded. Library size (*LS_j_*, number of UMIs in cell *j*) normalization and natural log transformation were applied to each cell library, i.e., 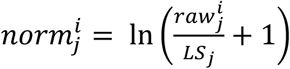, to quantify the expression of gene i in cell j, where 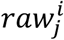 is the number of reads for gene *i* in cell *j*.

### Dimensionality reduction

Highly variable genes were defined as genes with greater than one standard deviation of the dispersion from the average expression of each gene. However, to account for heteroscedasticity, variable genes were identified separately in bins defined based on average expression. PCA analysis was performed on the normalized expression of the set of dataset-specific highly variable genes. We selected the top PCs based on gene permutation test (*74*). ‘Barnes-hut’ approximate version of t-SNE (*75*) (perplexity set to 30, 10k iterations) was applied on the top PCs to obtain a 2D projection of the data for visualization.

### Gene signature activation quantification

Gene signature activation was quantified relative to a technically similar background gene set as described in (*76*). Briefly, we identify the top 10 most similar (nearest neighbours) genes in terms of average expression and variance, then define the signature activation as the average expression of the signature genes minus the average expression of the background genes. GP66 tetramer staining signature definition is described in **Supplementary Note**. Additionally, we defined lists of ribosomal, mitochondrial, and cell cycle genes (*77*) for confounder controls (**Table S10**).

### High resolution clustering

Phenograph clustering (*39*) using the top PCs (see dimensionality reduction) was performed independently on each dataset to allow full control of the clustering resolution based on dataset-specific coverage and heterogeneity features. The clustering resolution (number of clusters) is controlled by the K nearest neighbour (KNN) parameter. We designed a simulation analysis to estimate the optimal clustering resolution, i.e., at what resolution the clustering is superior in quality to clustering driven by technical biases inherent to scRNAseq, as follows. Here we define the clustering quality as the clustering modularity reported by Phenograph, which indicates intra-cluster compactness and inter-cluster separation. The simulations consist of repeating the clustering analysis on 100 shuffled expression matrices to estimate the ‘null’ distribution of the clustering quality, where the gene expression measurements are permuted within each cell to retain the cell-specific coverage biases. We repeated this process for varying value of the KNN parameter *k* to compare the clustering modularity of the original *O*_k_ to the shuffled *S*_*k*_ data. The final resolution was defined as the maximal resolution where 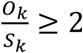 Pooled clustering analysis (joint rather than separated by dataset) and visualization was performed using PCA on the aggregate list of highly variable genes defined on each dataset. Clustering was done with and without controlling for confounding factors (number of UMIs, number of detected genes and gene signatures activation of ribosomal, mitochondrial, cell cycle and GP66 staining signature). Clustering analysis of TILs, dLN, and LCMV cells showed little overlap even after correcting for potential confounders.

After obtaining the initial clusters and identifying the overexpressed genes in each cluster, we apply two filters: (1) we exclude small clusters of B cells (CD79^+^ populations) from each dataset. (2) We identify PCs driven by B cell marker genes and remove the individual cells whose expression profile has high scores for those PCs (outliers). We then repeat the entire processing and clustering to prevent detecting highly variable genes and PCs driven by contaminations, which may in turn reduce the signal of other small populations of interest.

### Differential expression analysis and population matching

Differential expression was performed using Limma (version 3.32.10). We initially performed differential expression analysis between each cluster against the pool of all other clusters within a given dataset. Identified clusters were labelled as a known T cell subtype if the majority of the known subtype-defining genes were differentially over-expressed in that cluster. We then matched populations across experiments to assess the reproducibility of the populations and to uncover similarities across datasets that are masked due to overall tissue-context-specific differences. To reduce the effects of tissue-context-specific effects on the similarity calculation, we used the fold change (FC) measure of each gene *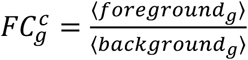* (average of gene *g* in cluster *c* (foreground) relative to all other clusters (background) of the same dataset). Then we measured the Pearson correlation between the FC vectors of all pairs of clusters across datasets. We compare this approach with an alternative approach that uses Euclidean distances between the average expression vectors, defined as average expression of all genes in a cluster and a recent data integration approach (*44*) following tutorial specifications [https://satijalab.org/seurat/immune_alignment.html; version 2.0.1].

### Robust cluster calling and robust population comparisons

For each dataset, we defined ‘robust clusters’ as those that had highly similar match in the biological replicate. High similarity is defined as Pearson correlation coefficient greater than *∼1.28* standard deviations from the mean for each dataset, corresponding to nominal p-value of 0.1. Hierarchical clustering was performed on the identified robust clusters using the inter-cluster similarity matrix, where the similarity was defined as above using the Pearson correlation between the FC vectors. Using the vector of average expression vectors did not achieve similar result; specifically, using hierarchical clustering of the Euclidean distances between the clusters average expression vector retained the grouping of clusters based on origin tissue (**Figure S3A**). We then analyzed differential expression patterns for clusters belonging to each meta-cluster, excluding cell cycle clusters. For a given pair of clusters of interest, A and B in datasets X and Y respectively, we performed three differential expression analyses: (1) differential expression in A relative to other clusters in X, (2) differential expression in B relative to other clusters in Y, and (3) differential expression in A relative to B. In addition to average expression differences, we quantified the detection rate of gene X as proportion of cells where 1 or more reads was mapped to X and prioritized differentially expressed genes exhibiting also differential detection across conditions. This analysis was performed for the two replicates separately and the results interpreted jointly; a gene was deemed as over-expressed in cluster A in tissue X if it is over-expressed relative to other clusters in X as well as relative to B, in both replicates.

### scRNAseq contour plots

Normalized scRNAseq expression measurements were visualized as contours, where zero (0) values were assigned random value drawn from a normal distribution centered around 0.

### Reversed Graph Embedding

Trajectory analysis of TIL populations (group I and II, excluding group III Tregs) was performed using Monocle (version 2.9.0, parameters max_components = 2, method = DDRTree).

### Gene signature definition

For each TIL subpopulation (group I Th1, group II Isc, group II nRes and group III Treg) we selected overexpressed genes exhibiting differential detection (as defined above) relative to all other TILs across both experiments (**Table S4**).

### Correspondence to human data

Human liver cancer TIL scRNAseq counts were downloaded from GEO [GSE98638]. Non-CD4^+^ T cells were filtered based on the classification in the original publication (*28*). Human gene symbols were translated to Mouse gene symbols using package biomaRt (version 2.37.8). Pre-processing, clustering and population matching analysis were applied as described above. Human melanoma TILs data scRNAseq counts were downloaded from GEO [GSE120575]. We selected CD4^+^ T cells as cells with at least one mapped read to CD4 and [CD3D or CD3E or CD3G], following the authors definition (*33*). 108 out of 136 Isc signature genes were mapped to human gene symbols. The detection rate of each Isc signature gene (as defined above) in each lesion were used to assess differential detection across responders and non-responders. We used two-sided Wilcoxon test to quantify the significance of differential activation.

### Correspondence with external gene signatures

Gene set enrichment analysis of immunologic gene signatures was performed using mSigDB (*61*) [C7: immunologic signatures database with clusterProfiler package (version 3.4.3). All other gene signatures were downloaded from the original publication’s supplementary materials. Correspondence to Tcmp signature was performed by differential expression of dLN *Ccr7*^+^ clusters n5-6 relative to other dLN and TIL (n1, n7-8, t1-7) rather than dLN subpopulations alone to satisfy the background conditions used in the original publication. The heterogeneity of the IL-27 co-inhibitory gene signature (*63*) was evaluated by analyzing differential gene expression across Th1, Isc, and Treg TIL, indicating which genes are preferentially expressed in one subpopulation versus the others.

## Acknowledgments

We thank Melanie S. Vacchio for cell sorting; Mariah Balmaceno-Criss and Qi Xiao for animal genotyping; Jack Greiner for the MC38 cell line; the NIH tetramer facility for reagents; the CCR Flow Cytometry Core for expert assistance; Yasmine Belkaid and Avinash Bhandoola for thoughtful discussions; Jonathan Ashwell, John O’Shea, Nicholas Restifo, Eytan Ruppin and Xin Wang for critical reading of the manuscript; and David Goldstein, Mariam Malik and the NCI Office of Science and Technology Resources for their support. This work used the NIH High performance computing cluster and was supported by the Intramural Research Program of the National Cancer Institute, Center for Cancer Research, National Institutes of Health.

## Author contributions

A.M., J.N., T.C., S.H. and R.B. designed research; A.M. designed and developed computational (bioinformatic) pipelines; A.M. and J.N. and T.C. performed research and analyzed data. S.T. guided TIL isolation procedures. D.M. provided advice on LCMV biology and LCMV viral stocks. M.M., Y. Z., and B.T. contributed to scRNA-seq capture. A.M. and R.B. wrote the manuscript with contributions from J.N. and S.H. S.H. and R.B. supervised the research.

## Competing interests

The authors declare no competing interests.

## Data and code availability

Data was deposited in [GEO GSE124691]. The computational pipeline is available on [https://github.com/asmagen/MagenSingleCell]. The pipeline requires access to Slurm high-performance computing core for efficient simulation analyses.

---

**Fig. S1:**
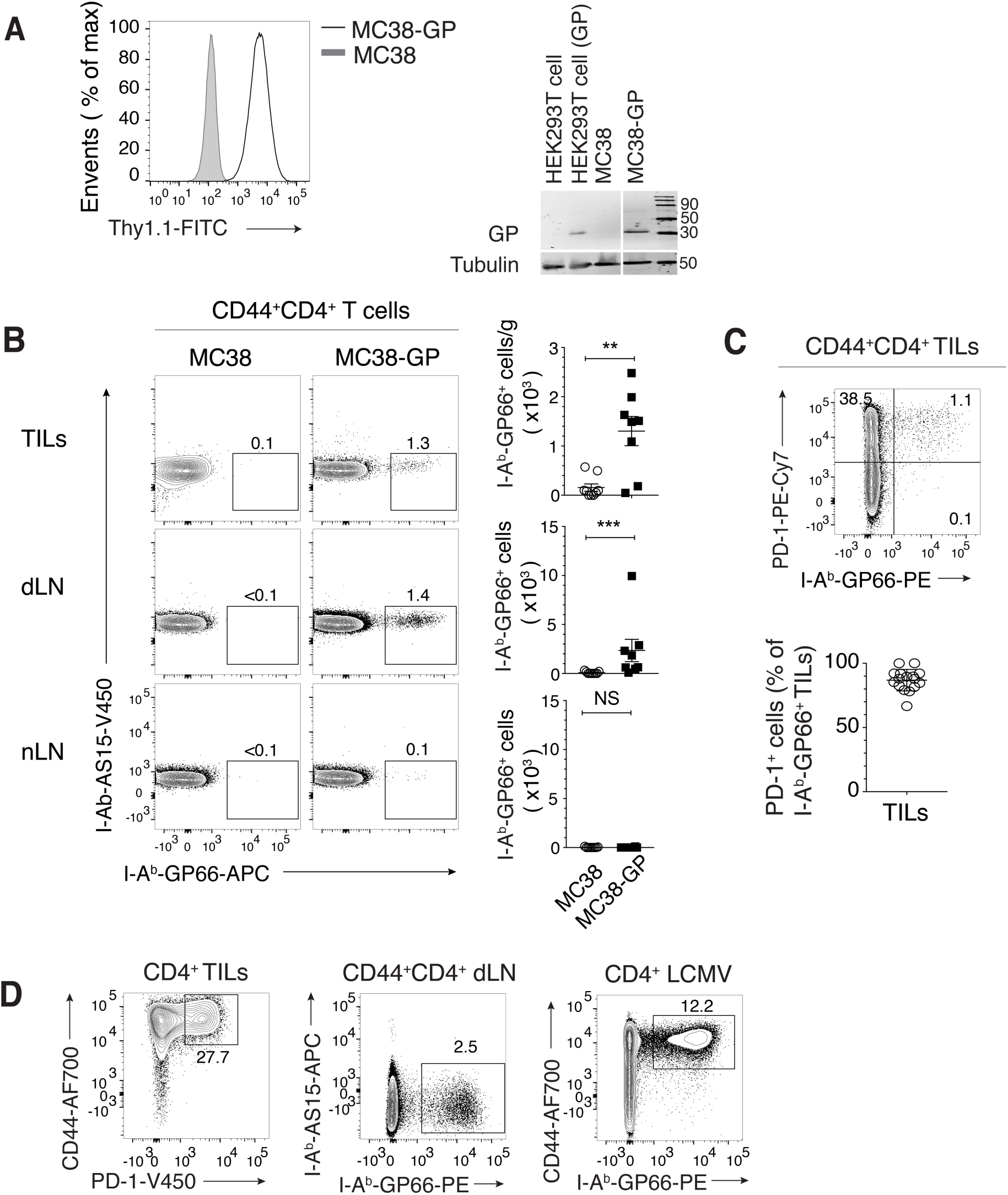
Characterization of antigen-specific CD4^+^ T cell responses in MC38 colon adenocarcinoma tumors. **(A)** Left panel shows overlaid protein expression of Thy1.1 in MC38 and MC38-GP cells. Right panel shows immunoblot analysis of GP protein expression in HEK293T cells, HEK293T cells transfected with pMRX-GP-IRES-Thy1.1 plasmid, MC38 cells or MC38-GP cells. **(B)** C57BL/6 mice were subcutaneously injected MC38 or MC38-GP cells and analyzed at day 14 post-injection. Left panel shows flow cytometry contour plots of GP66 vs. control (AS15 peptide from *T. gondii*) class II tetramer staining in TILs, dLN and nLN from MC38 and MC38-GP tumor-bearing mice. Right panel shows the number of GP66^+^ TILs per gram of tumor and total number of GP66^+^ dLN and nLN cells, separately for MC38 and MC38-GP tumor-bearing mice (Unpaired Mann-Whitney U test, * p < 0.01, ** p < 0.001, NS: not significant). **(C)** Top panel shows flow cytometry contour plots of GP66 tetramer staining vs. PD-1 in TILs. Bottom panel shows the percentage of PD-1^+^ cells out of GP66^+^ TILs. **(D)** GP66-specific CD44^hi^ CD4^+^ splenocytes were isolated from WT animals 7 days post-infection with LCMV Armstrong. Protein expression contour of populations used for scRNAseq captures from MC38-GP tumor-bearing mice (left: TILs PD-1 vs. CD44, middle: dLN GP66 vs. AS15 control) and LCMV Armstrong infected mice (right: GP66 vs. CD44).

**Fig. S2:**
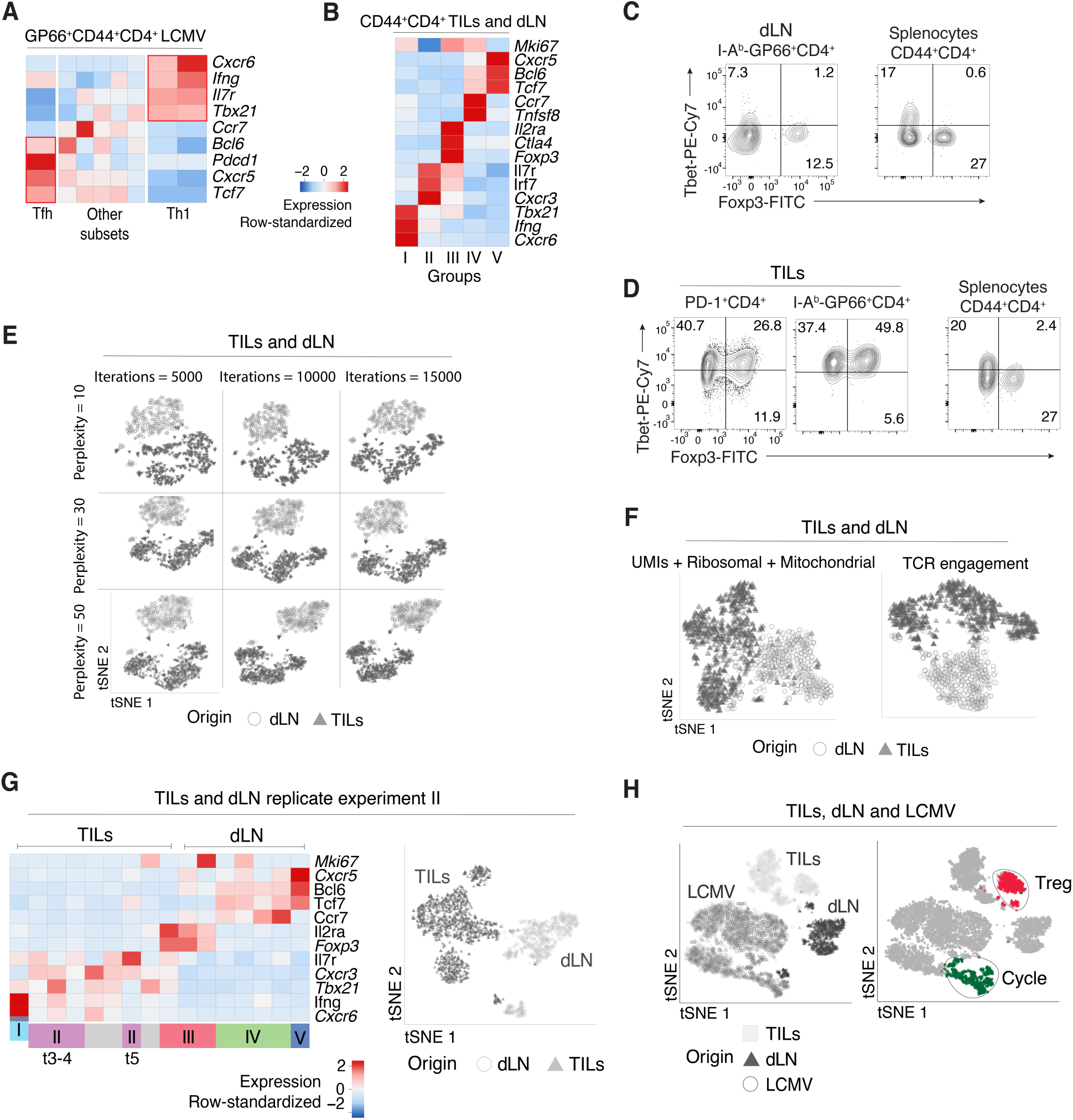
Characterization of immune responses to LCMV and MC38-GP by scRNAseq. **(A)** GP66-specific CD4^+^ splenocytes from WT animals 7 days post-infection with LCMV Armstrong analyzed by scRNAseq. Heatmap shows row-standardized expression of selected genes across LCMV clusters. **(B-G)** TILs and dLN cells from WT mice at day 14 post MC38-GP injection analyzed by scRNAseq. **(B)** Heatmap shows row-standardized expression of selected genes across main TIL and dLN groups (as defined in text). **(C)** Flow cytometry contour plots of Foxp3 vs. Tbet in CD44^hi^ GP66^+^ dLN cells (**left)** and in CD44^hi^CD4^+^ splenocytes from tumor-free mice control (**right**). **(D)** Flow cytometry contour plots of Foxp3 vs. Tbet in PD-1^+^ and GP66^+^ TILs (**left)** and in CD44^hi^ CD4^+^ splenocytes from tumor-free mice control (**right**). **(E)** tSNE display of TILs and dLN cells generated using different parameter combination of perplexity and number of iterations, grey-shaded by tissue origin. **(F)** tSNE displays of TILs and dLN cells, grey-shaded by tissue origin, post confounder correction for number of unique molecular identifiers (UMIs) and expression of ribosomal and mitochondrial coding genes (**left**) or TCR engagement on dLN cells as a result of GP66-tetramer-based purification (**right**). **(G)** scRNAseq analysis of TILs and dLN cells from replicate experiment II. Heatmap shows row-standardized expression of selected genes across TIL and dLN clusters (**left**). tSNE display of TILs and dLN cells, grey-shaded by tissue origin (**right**). **(H)** TILs, dLN and LCMV cells from replicate experiments I and II analyzed by scRNAseq. tSNE plots show TILs, dLN, and LCMV cells, grey-shaded by origin (**left**) or color-coded by Treg or cell-cycle (Cycle) clustering assignment (grey for all other clusters) (**right**).

**Fig. S3:**
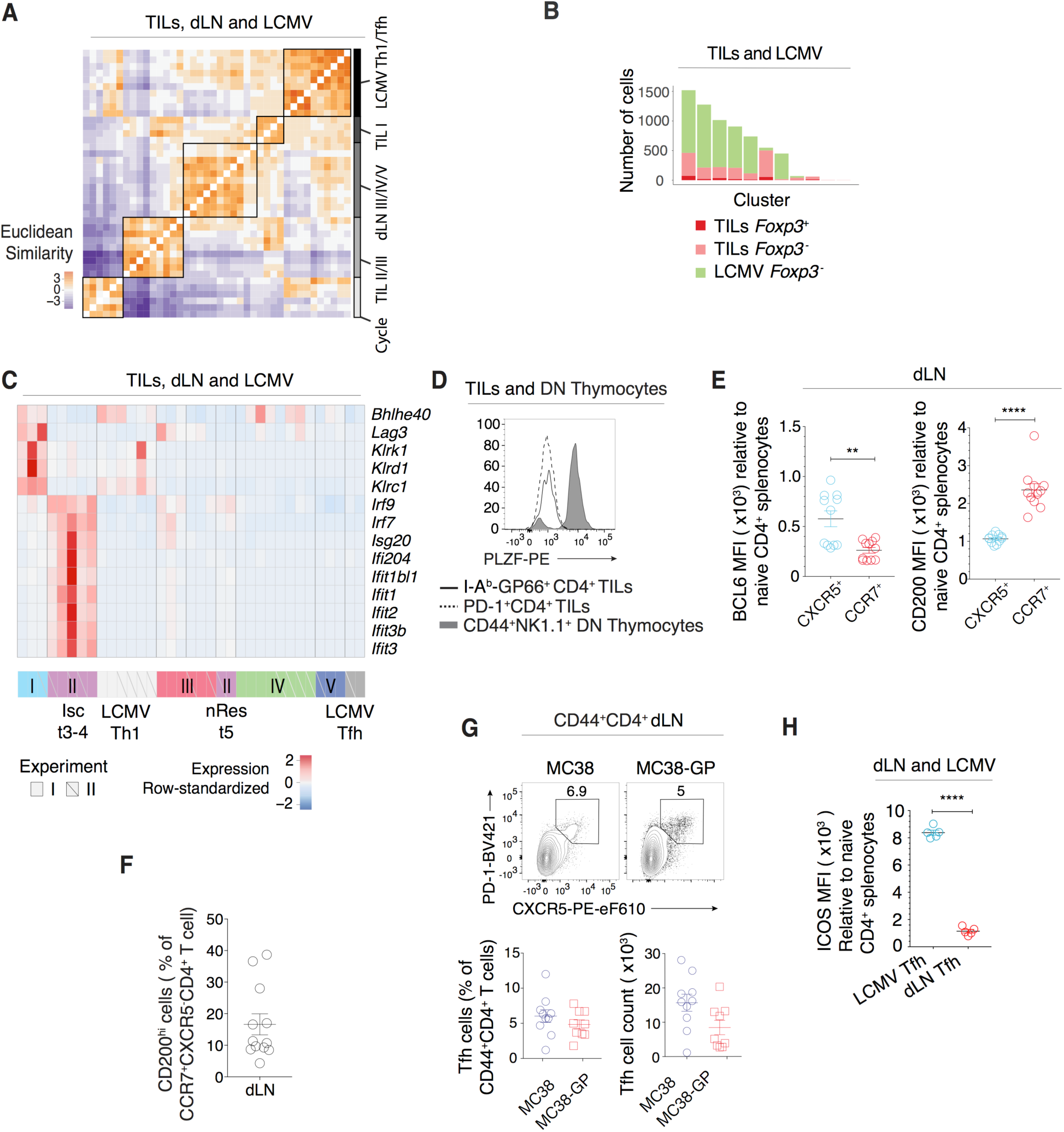
Assessment of tissue-context-specific effects on clustering analyses and TILs-dLN heterogeneity. **(A-C)** TILs, dLN and LCMV cells from replicate experiments I and II analyzed by scRNAseq. **(A)** Heatmap shows Euclidean similarity between cluster-specific average expression vectors (as defined in text) (**left**) annotated with cluster origin and cluster group or type (**right**). **(B)** Bar plot shows relative cluster composition of Foxp3^+^ or Foxp3^-^ TILs and Foxp3^-^ LCMV (no Foxp3^+^ cells found in GP66^+^ LCMV) after applying a data integration approach (*44*). **(C)** Heatmap shows row-standardized expression of TIL Isc and Th1 characteristic genes across TIL, dLN and LCMV clusters. **(D)** Overlaid protein expression of PLZF in GP66^+^ and PD-1^+^ TILs and CD44^hi^ NK1.1^+^ DN (double negative CD4^-^CD8^-^) thymocytes from tumor-free control mice. **(E)** Mean fluorescence intensity (MFI) of BCL6 and CD200 in CXCR5^+^ or CCR7^+^GP66^+^ dLN cells relative to naive CD4^+^ splenocytes from tumor-free control mice (Unpaired t-test, ** p < 0.005, **** p < 0.0001). **(F)** Percentage of CD200^hi^ cells out of CCR7^+^CXCR5^+^ dLN cells. **(G)** Top panel shows flow cytometry contour plots of CXCR5 vs. PD-1 in CD44^hi^ CD4^+^ dLN cells from MC38 and MC38-GP tumor-bearing mice. Bottom panel shows percentage of Tfh cells out of total CD44^hi^ CD4^+^ T cells in dLN (**left**) and total number of Tfh cells (**right**). **(H)** Mean fluorescence intensity (MFI) levels of ICOS in LCMV Tfh and dLN Tfh relative to naive CD4^+^ splenocytes from tumor-free control mice (Unpaired t-test, p < 10^−5^).

**Fig. S4:**
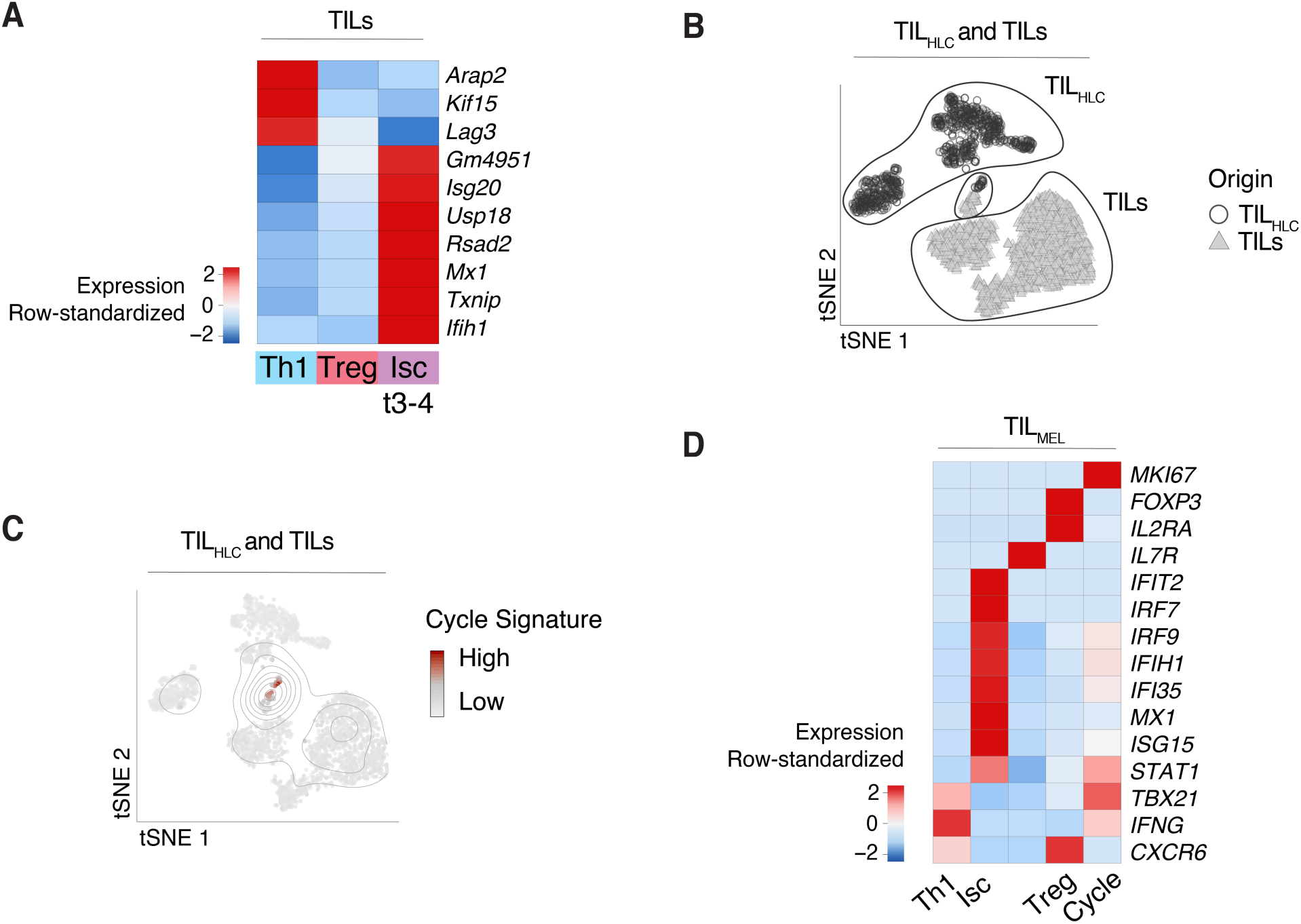
Correspondence to human data and dysfunction gene signatures. **(A)** Heatmap shows row-standardized expression of selected exhaustion genes across TIL Th1, Treg and Isc clusters (respectively clusters t1-2, t6-7 and t3-4 as shown in Fig. 1A). **(B-C)** Analysis of TIL_HLC_ and TILs (as defined in text). **(B)** tSNE plots show cells grey-shaded by origin. **(C)** tSNE plots show cells color-coded by cell cycle signature activation level. **(D)** Analysis of TIL_Mel_ (as defined in text). Heatmap shows row-standardized expression of selected TIL characteristic genes across TIL_Mel_ clusters.

## Supplementary Note

GP66-tetramer binding results in potential cross-linking of and signaling by the TCR of GP66-specific T cells. To model the transcriptomic effect of TCR engagement as a result of GP66-tetramer-based purification, we sought to compare LCMV-specific CD4^+^ T cells obtained either after GP66-tetramer purifcation or without tetramer-based purification. To enrich in such cells without tetramer staining, we noted that ∼94% of GP66-specific CD4^+^ splenocytes from LCMV-infected mice express little or no IL7R [IL-7 receptor *α* chain] (**Suppl. Note Figure A**). Thus, we considered that most CD44^hi^CD4^+^Il7R^+^ splenocytes were not LCMV-specific, and sorted CD44^hi^ IL7R^−^ (LCMV IL7R^−^) T cells for scRNAseq; in addition to antigen-specific CD44^hi^ GP66-tetramer purified (LCMV GP66^+^) T cells (**Suppl. Note Figure B**). Pooled clustering of the two samples revealed 2 (out of 6) clusters heavily dominated by stained cells (**Suppl. Note Figure C, top**), suggesting staining bias limited to those clusters. As expected from GP66 tetramer engagement with the TCR, GP66-specific clusters were characterized by genes involved in T cell receptor signaling and NFKB signaling (**Table S9**), while clusters containing cells from both samples displayed features of Tfh and Th1 cells (**Suppl. Note Figure C, bottom**). We designated the GP66-characteristic genes as the TCR engagement GP66 signature (**Table S10**) and regressed the activation scores of the signature from the expression matrix using a linear regression model fitted to each gene.

**Supplementary Note Figure:**
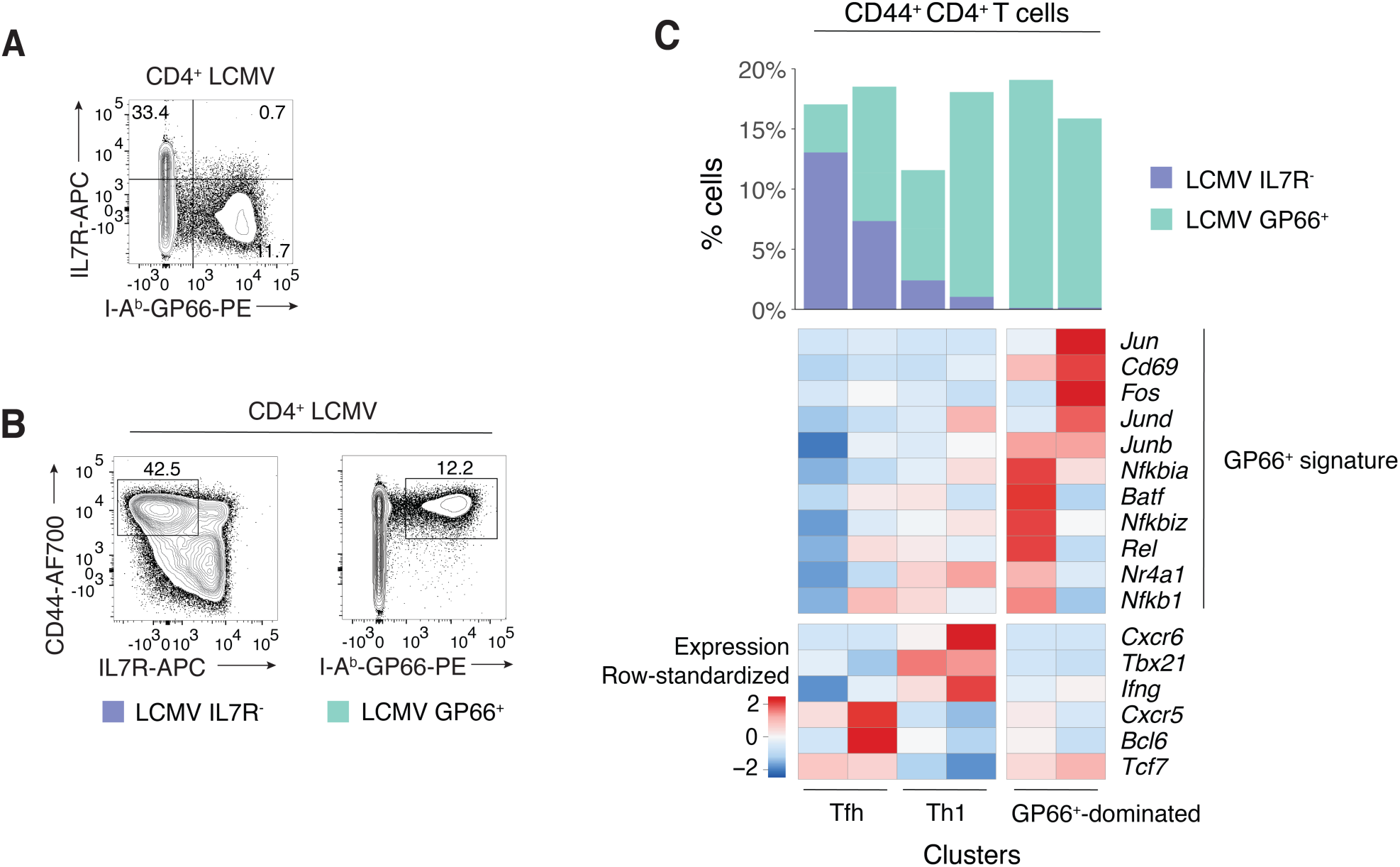
Transcriptomic effects of TCR engagement as a result of GP66-tetramer-based purification. **(A-B)** Analysis of CD4^+^ splenocytes from C57BL/6 animals 7 days post-infection with LCMV Armstrong. **(A)** Flow cytometry contour plot of GP66 tetramer staining vs. IL7R in CD4^+^ LCMV cells. **(B)** Flow cytometry contour plots of IL7R vs. CD44 (for LCMV IL7R^-^ sample, **left**) and GP66 vs. CD44 (for LCMV GP66^+^ sample, **right**). **(C)** LCMV IL7R^-^ and LCMV GP66^+^ cells analyzed by scRNAseq. Heatmap shows row-standardized expression of selected genes across pooled LCMV IL7R^-^ and LCMV GP66^+^ clusters (**bottom**). Bar plot indicates the number of LCMV IL7R^-^ and LCMV GP66^+^ cells in each cluster relative to the total number of cells (**top**).

